# Interindividual variations in peak alpha frequency do not predict the magnitude or extent of secondary hyperalgesia induced by high-frequency stimulation

**DOI:** 10.1101/2025.06.02.657417

**Authors:** Louisien Lebrun, Gloria Ricci, Arthur Courtin, Emanuel N. van den Broeke, Cédric Lenoir, André Mouraux

**Author notes:** CORRESPONDING AUTHOR, ADDITIONAL CO-CORESPONDING AUTHOR.

## Abstract

**Background:** Previous studies have shown an association between interindividual variations in the frequency of alpha-band EEG oscillations such as estimates of peak alpha frequency (PAF) and pain sensitivity. Whether differences in PAF also influence the susceptibility to develop central sensitization (CS) is unknown.

**Objective:** This study aimed to determine if the PAF of vision- and sensorimotor-related alpha-band activity is associated with the magnitude and extent of secondary mechanical hyperalgesia induced by high-frequency stimulation (HFS) of the skin, a surrogate marker of CS.

**Methods:** The EEG was recorded in 32 healthy participants at rest during eyes open and eyes closed conditions, and during bilateral finger movements. Two methods were used to isolate vision- and sensorimotor-related alpha-band activity based on sensitivity to eye closure and movement: one based on an independent component analysis, the other on spectral subtraction. PAF was assessed using a center-of-gravity approach or the Fitting Oscillations and One-Over-F (FOOOF) method, accounting for the aperiodic EEG component. HFS was applied to the right forearm, and pinprick sensitivity was assessed at both forearms, before and 40 minutes after HFS.

**Results:** Neither sensorimotor-nor vision-related PAF were significantly correlated with the magnitude or extent of HFS-induced secondary hyperalgesia. Interestingly, at the non-sensitized forearm, participants with a higher vision-related PAF exhibited greater habituation to pinprick stimuli, suggesting that variations in PAF may relate to variations in sensory habituation.

**Conclusion:** Interindividual variations of PAF were not significantly associated with the susceptibility to develop HFS-induced secondary hyperalgesia.

**New and Noteworthy:** Using several methods to estimate vision- and sensorimotor-related peak alpha frequency (PAF) in the EEG frequency spectrum, we found no significant association between interindividual variations in PAF and the susceptibility to develop secondary hyperalgesia following high-frequency stimulation (HFS) of the skin in healthy participants.

## INTRODUCTION

Alpha-band oscillatory activity (8-12 Hz) is the most prominent feature of the EEG recorded during rest in awake humans. The alpha rhythm was first described by Hans Berger, nearly a century ago (1). A distinctive feature of that rhythm is its predominance over posterior areas of the brain, and the fact that its magnitude markedly increases when participants close their eyes. The sensitivity to eye closure of posterior alpha oscillations has led most authors to consider that this oscillatory activity reflects a default state of brain areas “resting” or “idling” in the absence of input (2). However, alpha-band oscillations are not restricted to the visual cortex. Gastaut et al. (1952) described the Rolandic mu rhythm oscillating between 8-12 Hz over the scalp vertex and bilateral sensorimotor areas (3). Present at rest, the mu rhythm decreases during movement execution (3–5), most pronouncedly in the hemisphere contralateral to the executed movement (5). It is also suppressed during motor imagery (6,7), modulated by mental effort (8), attention and fatigue (9), and relatively unaffected by opening/closing of the eyes (10). Studies comparing the effects of upper and lower limb movements found that the movement-related suppression of the mu rhythm follows the mototopic and somatotopic organization of primary motor (M1) and somatosensory (S1) cortices (5).

Several studies have reported that the frequency at which the EEG spectrogram peaks within the alpha-band, referred to as the “peak alpha frequency” (PAF), tends to be slower in patients with chronic pain (11–16). Other studies have found that a slower PAF, recorded in healthy participants before they experienced experimental pain is associated with a greater sensitivity to phasic heat pain (17), tonic pain produced by topical capsaicin (18) and deep tissue pain produced by nerve growth factor injection (19,20), whereas an opposite trend has been observed in a previous study with tonic heat pain (17). The PAF has been shown to vary across individuals, but to remain relatively stable over time (17,20,22), suggesting that PAF may reflect an individual trait somehow related to pain sensitivity.

An important property of the nociceptive system is its propensity to sensitize when exposed to repeated nociceptive input or following tissue injury (23–27). Sensitization of the nociceptive system involves changes at the level of peripheral nociceptors leading to an increased sensitivity to noxious stimuli (peripheral sensitization) (28) and changes at the level of the central nervous system leading to an increased synaptic transmission of nociceptive input (central sensitization) described extensively at the level of the dorsal horn in animal studies (29,30), but most likely also involving higher-order synapses (31). A perceptual consequence of sensitization is hyperalgesia, i.e., an increased sensitivity to pain (32). Following skin injury, hyperalgesia develops both within the injured area (primary hyperalgesia) and in the surrounding uninjured skin (secondary hyperalgesia) (33,34). While both peripheral and central sensitization may contribute to the development of primary hyperalgesia, secondary hyperalgesia is thought to result exclusively from central sensitization (34,35). A hallmark feature of secondary hyperalgesia is a prominent increase in sensitivity to mechano-nociceptive input as compared to thermo-nociceptive input, enabling its characterization by testing sensitivity to mechanical pinprick stimulation of the skin (36,37).

In a recent study conducted in patients scheduled for lateral thoracotomy, we found that interindividual variations in the pre-operative susceptibility to develop secondary hyperalgesia following high-frequency electrical stimulation of the skin (HFS) – a procedure used to induce central sensitization experimentally – was associated with a greater likelihood post-thoracotomy pain at 2 months (38). The primary aim of the present study was to test whether interindividual variations in PAF may be related to variations in this susceptibility to develop HFS-induced secondary hyperalgesia.

Supporting this hypothesis, several studies have suggested that non-invasive brain stimulation of the sensorimotor cortex can reduce pain sensitivity (39,40), and that this effect may be mediated through a modulation of M1-connected brain structures involved in descending pain modulation (41). Therefore, if variations of sensorimotor-related alpha band activities index variations in M1 function, interindividual differences in sensorimotor PAF could be associated with changes in the state of descending pain modulation and the susceptibility to develop central sensitization. Finally, since most previous reports of a relationship between PAF and sensitivity to pain have used models of sustained pain which are known to induce some amount of sensitization, these observations might be explained in part by a relationship between PAF and susceptibility to sensitize.

To test this hypothesis, the EEG was recorded in healthy participants with their eyes open at rest, their eyes closed at rest, and with their eyes open while performing repetitive bilateral finger movements. These three EEG recording conditions were used to functionally isolate, vision-related alpha-band oscillations whose magnitude is known to increase following closure of the eyes and sensorimotor-related alpha-band oscillations whose magnitude reduces during finger movements. As in a multiverse analysis (42), several methods were used to (1) isolate vision- and sensorimotor-related alpha-band activity and (2) estimate PAF. After the EEG recording, central sensitization was induced in each participant using HFS delivered to the right volar forearm skin. The magnitude and extent of HFS-induced secondary hyperalgesia was estimated by assessing the sensitivity to mechanical pinprick stimuli delivered to the sensitized and non-sensitized forearms before and 40 minutes after HFS. The collected data allowed us to examine whether interindividual variations in vision- and/or sensorimotor-related PAF predicts the susceptibility to develop HFS-induced secondary mechanical hyperalgesia.

## METHODS

### Participants

Descriptive statistics are presented as mean ± standard deviation (SD), unless otherwise specified.

A total of 32 right-handed healthy volunteers (16 women and 16 men aged 25.3 ± 4.5 years [range: 19-38]) were included in this study between June and July 2020. Hand dominance was confirmed using the Flinders Handedness Survey (43). The study was approved by the local ethics committee (Comité d’Ethique Hospitalo-Facultaire des Cliniques universitaires Saint-Luc-UCLouvain; B403201316436). All participants provided written informed consent and received financial compensation. The experiment was conducted in agreement with the Declaration of Helsinki. The decision to include 32 participants was guided by the correlation coefficients reported in previous studies investigating the relationship between PAF and pain ratings in healthy volunteers (r = 0.74 in (21) and r = 0.5 in (18). Based on these two studies, the estimated required sample sizes range from 12 to 38 participants to achieve a power of 0.90 with an alpha level of 0.05 (44).

Inclusion criteria were being healthy and aged 18-40 years. Exclusion criteria were (1) any history of cardiac or neurological pathology, (2) regular practice of activities involving repeated stimulation of the volar forearm skin (e.g., volleyball and handball players), (3) a dermatological condition involving the forearm, (4) a history of traumatic injury of the upper limb and (5) regular tobacco consumption, (6) having taken part in a previous experiment involving stimulation of the volar forearm(s). Participants were asked to sleep a minimum of 8 hours the night before the experiment, to refrain from using recreational drugs, psychotropics, analgesics, or other medications in the days leading up to the study.

### Experimental design

The time course of the experiment is summarized in Figure 1A. Participants were seated on a comfortable chair with their arms resting on a desk, volar forearms facing upwards. Secondary hyperalgesia was induced at the right volar forearm using HFS regardless of hand dominance. EEG recordings were performed 25 minutes before HFS (T-25), 5 minutes before HFS (T-5), one minute after HFS (T+1) and approximatively 35 minutes after HFS (T+35). Each EEG recording lasted 5 minutes and included three conditions: eyes open at rest, eyes closed at rest and eyes open while performing repetitive bilateral flexion-extension movements with their index fingers. During the 30 minutes pause separating T+1 and T+35 EEG recordings, participants were instructed to read a book or magazine of their choice. Sensitivity to mechanical pinprick stimuli was evaluated at both volar forearms 20 minutes before (T-20) and approximatively 40 minutes after (T+40) HFS. The extent of the HFS-induced area of secondary hyperalgesia was evaluated 40 minutes after HFS (T+40), at the HFS-treated arm.

**Figure 1.**
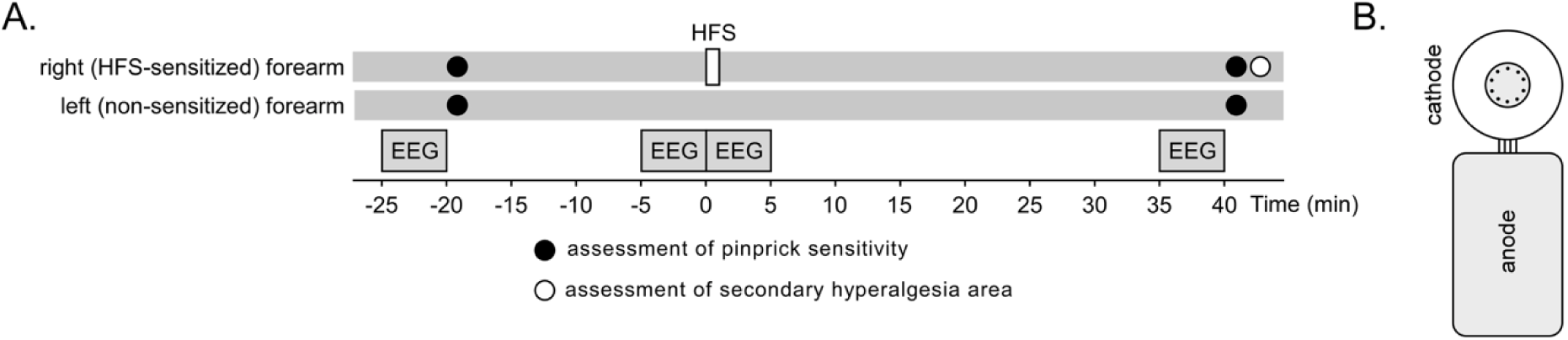
**A.** Timeline of the experiment. HFS was applied to the right volar forearm. EEG segments lasting 5 minutes and including three conditions (eyes open at rest, eyes closed at rest, eyes open while performing bilateral finger movements) were recorded 25 and 5 minutes before HFS (T-25 and T-5), immediately after HFS, and approximately 35 minutes after HFS (T+35). Pinprick sensitivity of the left and right volar forearms was assessed before HFS (T-20) and approximately 40 minutes after HFS (T+40). The spatial extent of the HFS-induced area of secondary hyperalgesia was assessed approximately 40 minutes after HFS (T+40). **B.** HFS was delivered to the forearm skin using a cathode consisting of 10 blunt tungsten pins arranged on a circle with a diameter of 5 mm and a 24×20 mm anode conductive gel pad placed next to the cathode.

### Induction of secondary hyperalgesia using HFS

HFS was applied to the right volar forearm, 10 cm distal from the cubital fossa using an electrode designed to preferentially activate cutaneous free nerve endings (EPS-P10, MRC Systems GmbH, Heidelberg, Germany). The electrode consisted of a multi-pin cathode and a flat large-surface anode (Figure 1B). The cathode consisted of 10 blunt tungsten pins arranged on a circle with a diameter of 5 mm. Each pin had a diameter of 0.25 mm and protruded by 0.65 mm over the base of the electrode. The anode was a 24 x 20 mm conductive gel pad placed next to the cathode. The stimulation consisted of 5 trains of 100 Hz charge-compensated biphasic electrical pulses (45) delivered using a constant-current electrical stimulator (DS5; Digitimer, Welwyn Garden City, United Kingdom) triggered by a National Instruments digital-analogue interface (NI6343, National Instruments, Austin, Texas, USA). Each train lasted 1 second (100 pulses/train). The time interval between the onset of each train was 10 s. The intensity of stimulation was set to 20 the detection threshold to a single pulse, which was estimated in each participant at the beginning of the experiment using the method of limits (0.126 ± 0.045 mA).

### Assessment of mechanical pinprick sensitivity at the volar forearms

Before HFS (T-20) and after HFS (T+40), the sensitivity to mechanical pinprick stimuli was assessed at the left and right volar forearms using a custom-made pinprick stimulator (0.25 mm probe diameter, CATL, UCLouvain, Belgium) exerting a 128 mN force (26,27,45). At each time-point, three pinprick stimuli were applied perpendicularly to the skin surrounding the area of HFS stimulation at the right volar forearm and on the same location at the left volar forearm. After each stimulus, participants were requested to report the intensity of the elicited sensation using a numerical rating scale ranging from 0 (no perception) to 100 (maximal imaginable pain), with 50 marking the limit between non-painful and painful sensations as used in several previous experiments (46–49). To avoid sensitization of the skin by repeated mechanical pinprick stimulation, the probe was not applied twice on the same location. The order of the testing (HFS-treated arm first or control arm tested first) was counterbalanced across participants and remained constant across the two time points. The mean of the three ratings obtained at each time point and arm was used as an index of pinprick.

### Assessment of the area of HFS-induced secondary hyperalgesia

The same 128 mN pinprick probe was used to map, after HFS (T+40), the area of increased mechanical pinprick sensitivity surrounding the site of HFS stimulation at the HFS-treated volar forearm. The stimuli were applied along 4 axes and 8 directions separated by a 45-degrees angle and each originating from skin location corresponding to the center of the HFS electrode and. Participants were asked to close their eyes during the mapping procedure. Starting away from the HFS-treated skin and moving towards the center, stimuli were applied every 5 mm. Participants were asked to report when the sensation increased, and the corresponding stimulation location was marked with a felt-tip pen (26,47,49–51). For each of the 8 directions, the distance between the center of the HFS electrode location and the mark was measured, and the area of secondary hyperalgesia was computed as the area of a piecewise polynomial fit of a 2D closed curve linking the 8 marked locations (‘interpclosed’ function; Santiago Benito 2023, Matlab Central File Exchange) (26,27). Since this function cannot process null values, we assigned a value of 0.1 when no secondary hyperalgesia area was present in this direction. The radius of the area of increased pinprick sensitivity was then computed as the square root of the estimated area divided by π, as if the area was in the shape of a circle. This transformation was performed prior to any analyses.

### EEG recording

The EEG was recorded using 64 scalp electrodes with active shielding, placed according to the International 10-20 system (ANT Neuro, Enschede, The Netherlands). The signals were digitized at 1,000 Hz using an average reference. All impedances were kept under 20 kΩ. At each time point (T-25, T-5, T+1 and T+36), the recording consisted of 3 minutes at rest with eyes open (participants were asked to relax and stare at a cross displayed 1 meter in front of them, at a 30° angle below eye level), followed by 1 minute at rest with eyes closed, and 1 minute with eyes open while performing repeated bilateral index finger movements at a rate of approximately 1 Hz while fixing their gaze on the cross. These three conditions (EYES-OPEN-REST, EYES-CLOSED-REST, EYES-OPEN-MVT) were used to differentially modulate vision-related alpha-band activity (which should be enhanced in the EYES-CLOSED-REST condition as compared to the other conditions) and sensorimotor-related alpha-band activity (which should be attenuated when performing finger movements in the EYES-OPEN-MVT condition as compared to the other conditions).

### EEG preprocessing

The EEG signals collected before HFS (T-25) were analyzed offline using the Letswave 6 toolbox (http://letswave.org) running on MATLAB 2018a (Mathworks, USA). The EEG data collected at the other time points was not analyzed in the present study.

First, the EEG recordings were cut in three segments: a 180-s EYES-OPEN-REST segment, a 60 s segment EYES-CLOSED-REST and a 60 s segment EYES-OPEN-MVT (6). The three segments were then filtered using a 0.5 - 47 Hz zero-phase Butterworth band-pass filter (order 4) and further segmented into half-overlapping 5-s epochs. Each segment was demeaned, and a linear detrend was applied to remove slow drifts of the signal. Noisy electrodes (1-2 electrodes in 10 out of 32 recordings) were interpolated using the average of the three nearest electrodes. Epochs with clear artifacts (absolute amplitude ≥200 μV) were rejected (2.4% of epochs from EYES-OPEN-REST condition, 2.8% of epochs from EYES-CLOSED-REST condition and 2.8% of epochs from EYES-OPEN-MVT condition). Signals were re-referenced to the average of all channels excluding the two mastoid electrodes (M1, M2).

### Functional separation of vision- and sensorimotor-related alpha-band activity

In previous studies, PAF has been measured using various methods (17,18,21,52–54) and at different scalp locations (17,55–59), with some studies interpreting it as a marker of sensorimotor activity even when recorded over central or occipital regions or when clearly responding to closure of the eyes (11,17–19). In this work, we aimed to explore multiple approaches to separate vision- and sensorimotor-related alpha-band activity and estimate their PAF.

### ICA-based method

In a first approach, an independent component analysis (ICA) was used to dissociate visual alpha-band activity (VISUAL-ALPHA) and sensorimotor alpha-band activity (SM-ALPHA) based on their sensitivity to closure of the eyes and movement of the hands.

For each participant, the ‘runica’ algorithm was used to separate the 62 EEG scalp signals into 61 independent components (ICs) having a fixed topographical distribution, and maximally independent time courses. In the case channels were interpolated during preprocessing, the number of ICs was lowered by the number of interpolated channels (60). Because the amplitude of VISUAL-ALPHA should be selectively increased during the EYES-CLOSED-REST condition while SM-ALPHA should be selectively decreased during bilateral moving of the fingers in the EYES-OPEN-MVT condition, ICA can be expected to capture visual- and sensorimotor-related alpha-band activity into separate ICs.

For each participant and each IC time course, a Fast-Fourier Transform (FFT) was used to compute the amplitude frequency spectra of all 5-s half-overlapping epochs, windowed using a Hanning function. The spectra were then averaged per participant. The obtained frequency spectra were used to identify ICs capturing alpha-band activity, and to categorize these ICs into the following three categories: VISUAL-ALPHA (ICs capturing vision-related alpha-band activity that was increased in the EYES-CLOSED-REST condition as compared to EYES-OPEN-REST condition and the EYES-OPEN-MVT condition), SM-ALPHA (ICs capturing sensorimotor-related alpha-band activity that was decreased in the EYES-OPEN-MVT condition as compared to the EYES-OPEN-REST condition and was not increased in the EYES-CLOSED-REST condition as compared to the EYES-OPEN-REST condition) and MIXED-ALPHA (ICs capturing alpha-band activity that was both sensitive to closure of the eyes – greater activity in the EYES-CLOSED-REST condition compared to the EYES-OPEN-REST condition – and sensitive to finger movements – reduced activity in the EYES-OPEN-MVT condition compared to the EYES-OPEN-REST condition).

The classification was achieved for each IC of each participant as follows. First, using the average of the frequency spectra obtained in all three conditions, the peak of potential alpha-band activity captured by the IC was computed using the center of gravity (CoG) formula (19,61)

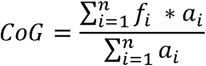

where f is the i^th^ frequency bin between 9 and 11 Hz, a is the spectral amplitude for f_i_, and n is the number of frequency bins between 9 and 11 Hz. This value was then used to define a 6 Hz frequency window of interest centered on the estimated CoG. Second, the mean amplitude and standard error of the mean (SEM: standard deviation of amplitudes divided by the square root of the number of bins) were computed separately for each of the three EEG recording conditions using all amplitude bins located within the selected window of interest. The mean and SEM were used to compute a confidence interval corresponding to the mean ± 1.97*SEM, and the lower and upper bounds of these confidence intervals (CI) were used to perform the categorization. ICs were classified as capturing vision-related alpha-band activity (VISUAL-ALPHA) when the lower bound of the CI in the EYES-CLOSED-REST condition exceeded the upper bound of the CI in the EYES-OPEN-REST condition (indicating an increase of alpha during eye closure), whereas the CI in the EYES-OPEN-REST condition overlapped the CI in the EYES-OPEN-MVT condition (indicating no strong effect of movement on alpha-band amplitude). Similarly, ICs were classified as capturing movement-related alpha (SM-ALPHA) when the lower bound of the CI in the EYES-OPEN-REST condition exceeded the upper bound of the CI in the EYES-OPEN-MVT condition (indicating a movement-induced reduction of alpha), whereas the CI in the EYES-OPEN-REST and EYES-CLOSED-REST conditions overlapped (indicating low sensitivity to closure of the eyes). Finally, ICs were classified as MIXED-ALPHA when they were both sensitive to closure of the eyes (lower bound of the CI in the EYES-CLOSED-REST condition > upper bound of the CI in the EYES-OPEN-REST condition) and sensitive to the finger movements (lower bound of the CI in the EYES-OPEN-REST condition > upper bound of the CI in the EYES-OPEN-MVT condition). Examples of ICs capturing vision-related and sensorimotor-related alpha-band activity are shown in Figure 2.

**Figure 2.**
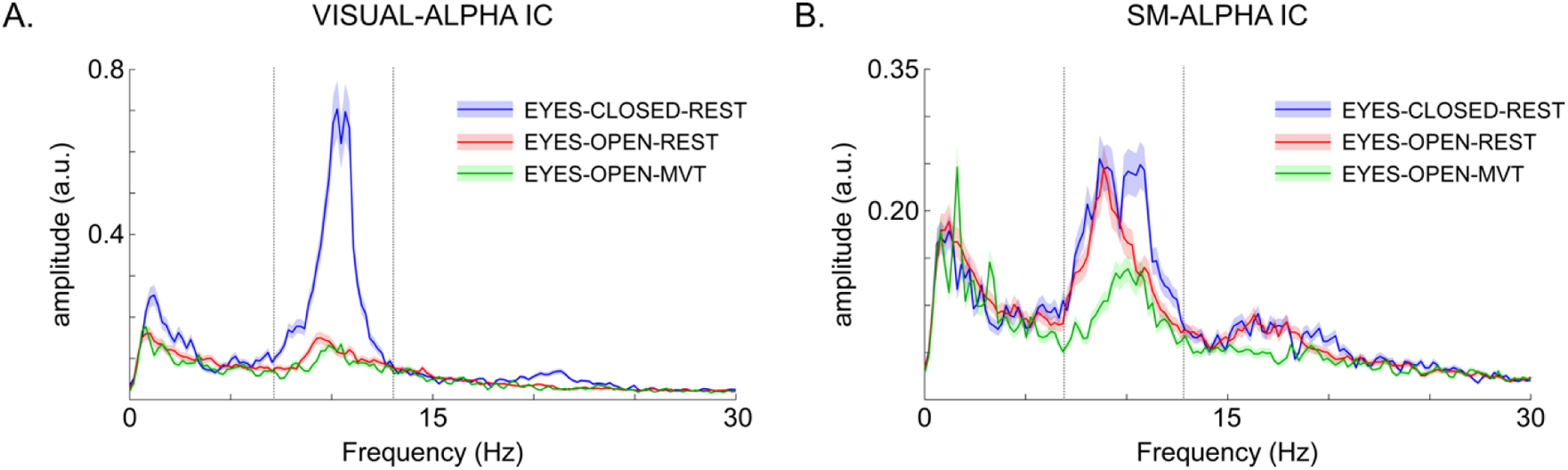
A. EEG frequency spectrum of a representative independent component (IC) categorized as capturing vision-related alpha-band activity (VIS-ALPHA) that was increased during closure of the eyes (EYES-CLOSED-REST condition) as compared to the eyes open conditions (EYES-OPEN-REST and EYES-OPEN-MOVEMENT). B. EEG frequency spectrum of a representative IC categorized as capturing sensorimotor-related alpha-band activity (SM-ALPHA) that was decreased during finger movements (EYES-OPEN-MVT condition) as compared to rest (EYES-OPEN-REST and EYES-CLOSED-REST conditions). Y-axis: arbitrary units. X-axis: frequency in Hz. The vertical grey lines represent the 6 Hz frequency window used to assess the alpha-band activity.

For each participant, the VISUAL-ALPHA, SM-ALPHA and MIXED-ALPHA ICs were then used to reconstruct three separate EEG datasets maximally capturing vision-related, sensorimotor-related and mixed alpha-band activity, respectively.

### Subtraction-based method

Suppression of movement-related activity in the EYES-OPEN-MVT condition may be expected to constitute the main difference between that condition and the EYES-OPEN-REST condition. Therefore, computing the difference between the EYES-OPEN-REST and EYES-OPEN-MVT spectra could constitute a simple alternative to efficiently isolate movement-related alpha-band activity. Similarly, because closure of the eyes selectively enhances vision-related alpha oscillations, subtraction of the EYES-OPEN-REST and EYES-CLOSED-REST spectra could constitute a means to isolate vision-related alpha-band activity. As compared to the ICA-based approach, a possible advantage of this alternative approach is that the subtraction can be expected to cancel out the contribution of activity that is equally present in both of the subtracted spectra, including the 1/f aperiodic component of the EEG power spectrum that can bias estimation of PAF (18).

### Estimation of peak alpha frequency (PAF)

Vision-related alpha oscillations can be expected to predominate over occipital electrodes, whereas sensorimotor-related alpha oscillations modulated by bilateral movements of the fingers can be expected to spread over left and right central electrodes (5,10). Therefore, the PAF of vision-related alpha-band activity was estimated in the average of the frequency spectra of occipital electrodes O1, Oz and O2, using the eyes-closed derived VISUAL-ALPHA spectra of the ICA-based method and the EYES-CLOSED-REST minus EYES-OPEN-REST spectra of the subtraction-based method. Similarly, the PAF of movement-related alpha-band activity was estimated in the average of the frequency spectra of central electrodes C3, Cz and C4, using the eyes open-derived SM-ALPHA spectra of the ICA-based method and the EYES-OPEN-REST minus EYES-OPEN-MVT spectra of the subtraction-based method. Finally, because the MIXED-ALPHA signals generated by the ICA-based method might have included both vision- and sensorimotor-related alpha-band activity, its PAF was estimated both at occipital and at central channels.

Two different methods were used to estimate the PAF: the center of gravity (CoG) method and the Fitting Oscillations and One Over F method (FOOOF (62)).

For the CoG method, we used the formula described above and a 9-11 Hz interval as was used in previous studies assessing the relationship between PAF and sensitivity to pain (18,19). As visual inspection of the EEG frequency spectra indicated that the PAF of some individuals was located outside the 9-11 Hz interval, the CoG was also estimated using a larger 7-13 Hz window. In the subtracted spectra, some amplitude values within the 9-11 or 7-13 Hz windows were negative. Only positive values were used for the CoG estimation.

For the FOOOF method, the spectrum was decomposed in its periodic and aperiodic components, as described in Donoghue et al. (62). In this case, the alpha band was defined as the range from 7 to 14 Hz and default algorithm parameters were employed, in coherence with other studies that performed similar analyses (63). When using this approach a larger frequency range is advised, as the overlapping of the frequency borders with a spectral peak is associated with higher parameter estimation errors (64). After the decomposition, only the periodic component was considered, as it is expected to contain alpha-band oscillatory activities. A gaussian curve was used to fit the periodic component of the spectrum, and its parameters were used to characterize the PAF (defined as the center of the gaussian curve). As the FOOOF method is intended to suppress the 1/f aperiodic component of the EEG, it was not applied to the subtracted spectra.

### Statistical analysis

Statistical analyses were performed using JASP (JASP Version 0.14.1.0 for Windows). Statistical tests were chosen depending on the distribution of the dependent variables, which was assessed by visual examination of the histograms for skewness and kurtosis, and the Q-Q plots of the residuals.

#### HFS-induced change in pinprick sensitivity

Mechanical pinprick ratings followed a close-to-normal distribution and we thus conducted a repeated-measures ANOVA with two within-subjects factors: TIME (before vs. after HFS) and ARM (HFS-treated vs. control arm), and one between-subjects factor: ORDER (whether the HFS-treated or control arm was tested first). Data were averaged per condition for each participant prior to analysis.

#### Peak-alpha frequency

A correlation analysis was used to evaluate the relationship between sensorimotor- and vision-related PAF and the changes in pinprick sensitivity after HFS (change in pinprick sensitivity at the sensitized HFS-treated forearm: ΔNRS_HFS_=NRS_HFS_(T+40) - NRS_HFS_(T-20), change in pinprick sensitivity at the non-sensitized forearm: ΔNRS_NS_=NRS_NS_(T+40) - NRS_NS_(T-20), difference in the change in pinprick sensitivity at the two forearms: ΔΔNRS = ΔNRS_HFS_ - ΔNRS_NS_), and between PAF and the radius of the area of secondary mechanical hyperalgesia (R). For each estimated PAF, the analyses were performed using Spearman’s correlations because normality was not strictly respected for all variables. Yet, as it was respected for our primary aim (relationship between sensorimotor PAF and secondary hyperalgesia), we displayed Pearson’s correlation for these analyses.

To examine a possible relationship between PAF and pain sensitivity, we also computed Spearman’s correlation between PAF and the detection threshold to single electrical stimuli, as well as the average ratings of the intensity of the sensation elicited by the HFS trains. Finally, we also assessed the correlation between inter-individual variations in vision- and sensorimotor-related PAF, and between the estimates of PAF obtained using the different approaches (ICA vs subtraction-based method to isolate vision- and sensorimotor-related alpha-band activity, CoG vs FOOOF method to estimate PAF).

Because the primary aim of our study was to test a possible association between PAF of sensorimotor alpha-band oscillations and the susceptibility to develop central sensitization, and because previous studies showed that the extent of the area of secondary hyperalgesia may be a more reliable readout as compared to changes in pinprick ratings (47), our primary outcome was the correlation between the sensorimotor PAF and the radius of the HFS-induced area of increased pinprick sensitivity. All other correlations were considered as exploratory, thus we did not apply corrections for multiple comparisons.

## RESULTS

### Pinprick sensitivity before and after HFS

After HFS, all but one participant reported an increased sensitivity to pinprick stimuli at the sensitized forearm as compared to the control arm (Figure 3A). The repeated-measures ANOVA showed a significant TIME × ARM interaction (F(1, 30) = 48.414, p = <.001, η²[z] = 0.617).

**Figure 3.**
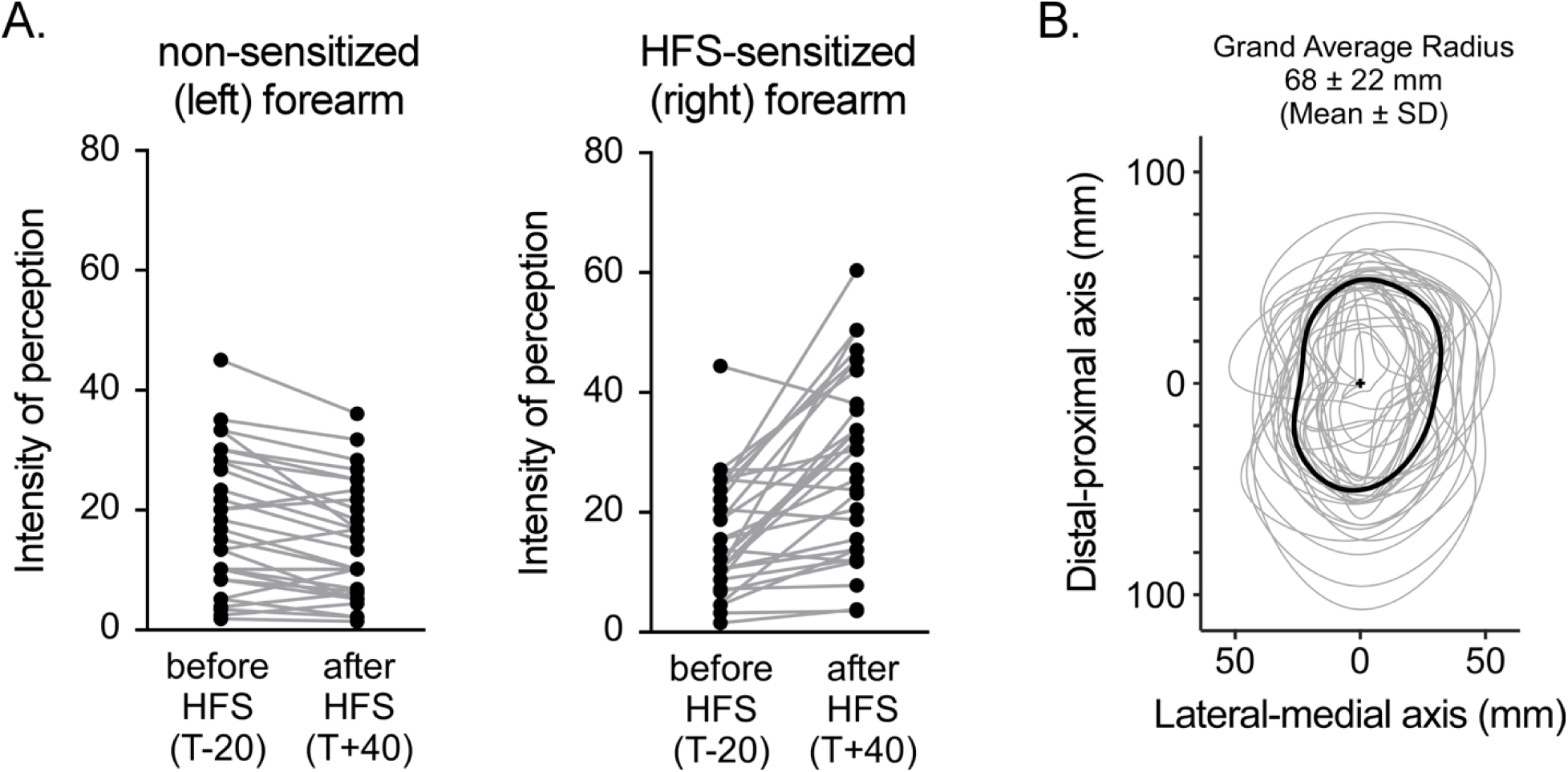
**A**. Individual intensity ratings for pinprick stimuli delivered to the HFS-sensitized forearm and the contralateral non-sensitized forearm, before and after HFS (T-20 and T+40). Note the clear increase in pinprick ratings at the sensitized forearm due to HFS-induced sensitization, and the decrease in pinprick ratings at the non-sensitized forearm possibly due to perceptual habituation. **B**. Estimated area of HFS-induced secondary hyperalgesia (grey contours: individual participants; black contour: group-level average). The origin (+) corresponds to the center of the cathode of the HFS electrode.

Post hoc comparisons confirmed a significant increase in pinprick ratings at the HFS-treated arm, with a mean increase of 11.854 (t(31) = 5.424, p < 0.001, Cohen’s d = 0.959 [0.533 to 1.374], 95% CI for mean difference: 7.397 to 16.311). Conversely, there was a significant decrease in ratings at the control arm, with a mean reduction of 3.312 (t(31) = –4.177, p < 0.001, Cohen’s d = –0.738 [–1.126 to –0.342], 95% CI for mean difference: –4.930 to –1.695).

### HFS-induced area of secondary hyperalgesia

At the HFS-treated forearm, an area of increased pinprick sensitivity was found in all but one participant (Figure 3B). Across participants showing an area of increased sensitivity, the extent was 101 ± 32 mm along the proximal-distal axis, 53 ± 20 mm along the lateral-medial axis, 77 ± 29 mm and 66 ± 27 mm along the two diagonal axes. The grand average estimated area radius was 68 ± 22 mm.

### Vision- and sensorimotor-related alpha-band activity isolated using the ICA-based method

After blind source separation using ICA, 19.4 ± 5.5 ICs capturing alpha-band activity were identified in 31 out of 32 participants, of which 11.9 ± 5.4 ICs were categorized as vision-related (VISUAL-ALPHA) in 31 out of 32 participants, 5.5 ± 5 ICs were categorized as sensorimotor-related (SM-ALPHA) in 24 out of 32 participants and 4.5 ± 4.3 ICs were categorized as mixed (MIXED-ALPHA) in 22 out of 32 participants.

The reconstructed VISUAL-ALPHA signals clearly captured vision-related alpha-band activity that increased during closure of the eyes, was only minimally affected by finger movements and, most importantly, displayed an occipital scalp topography compatible with activity originating predominantly from visual areas (Figure 4A). Similarly, the SM-ALPHA signals captured sensorimotor-related alpha-band activity that decreased during movement of the fingers, was minimally influenced by closure of the eyes, and exhibited a bilateral centroparietal scalp topography compatible with activity originating from the representation of the left and right hands within the sensorimotor cortex (Figure 4B). The reconstructed MIXED-ALPHA signal captured alpha-band activity that predominated over posterior regions and tended to both increase during closure of the eyes and decrease during finger movements (Figure S1).

**Figure 4.**
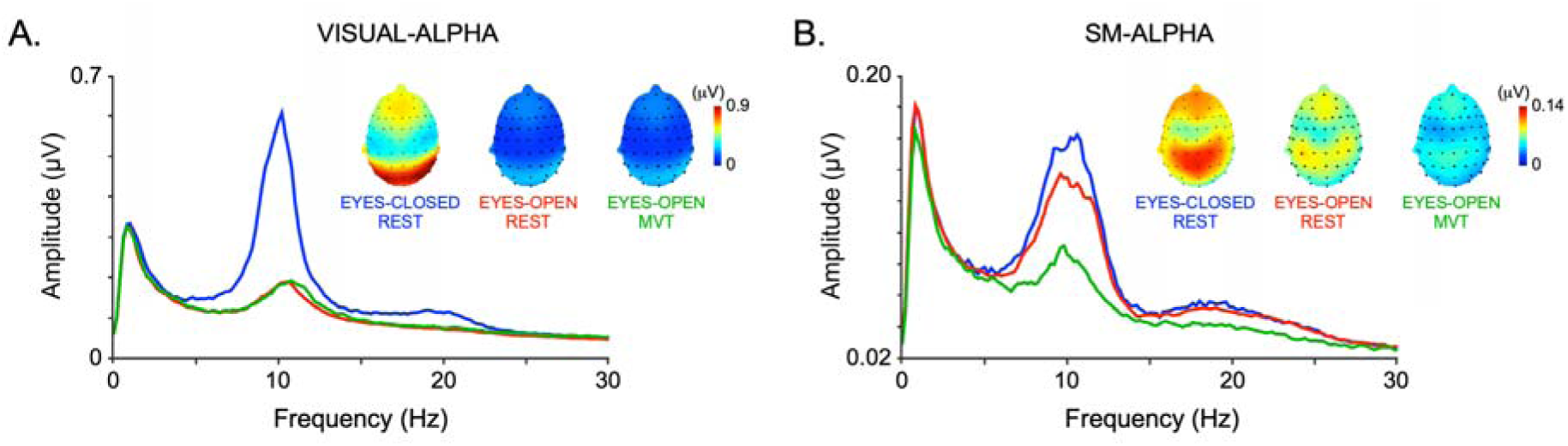
**A.** Group-level average frequency spectra of the ICA-separated EEG signals capturing vision-related alpha-band activity (VISUAL-ALPHA signals) at occipital electrodes in the EYES-CLOSED-REST (blue), EYES-OPEN-REST (red) and EYES-OPEN-MVT (green) conditions. Note the clear increase in alpha-band amplitude during closure of the eyes. Also note the occipital topography of the peak. **B.** Group-level average frequency spectra of the ICA-separated EEG signals capturing sensorimotor-related alpha-band activity (SM-ALPHA signals) at central electrodes. Note the selective decrease of amplitude during finger movements, and the bilateral central-parietal topography compatible with activity originating from the left and right sensorimotor cortices.

### Vision- and sensorimotor-related alpha-band activity isolated using the subtraction-based method

The group-level averages of the subtracted frequency spectra are shown in Figure 5. As expected, the subtraction spectrum EYES-CLOSED-REST minus EYES-OPEN-REST exhibited a clear peak of alpha-band activity that was maximal over occipital electrodes, compatible with alpha oscillations originating predominantly from the visual cortex (Figure 5A).

**Figure 5.**
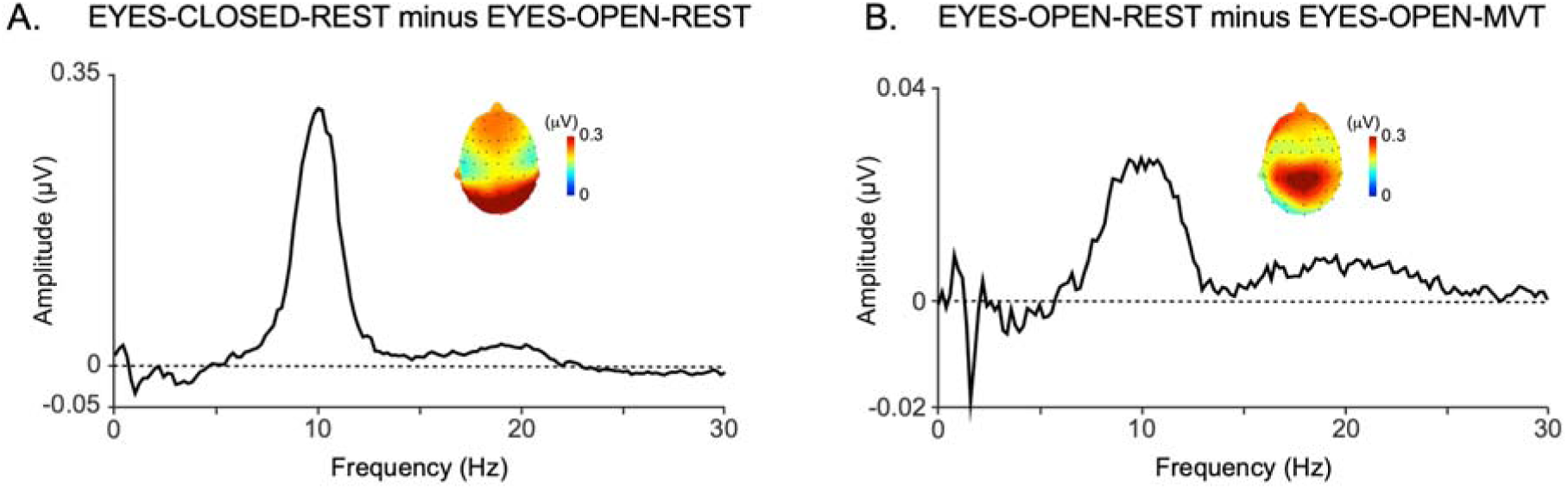
**A.** Group-level average of the subtracted EEG frequency spectra EYES-CLOSED-REST minus EYES-OPEN-REST conditions at occipital electrodes. Note the clear occipital topography of the peak indicating that this approach effectively isolated vision-related alpha-band activity. **B.** Group-level average of the subtracted frequency spectra EYES-OPEN-REST minus EYES-OPEN-MVT conditions at central electrodes. Note the bilateral central topography of the peak indicating that this approach succeeded in isolating sensorimotor-related alpha-band activity. Also note the lack of an 1/f decay (i.e. the aperiodic component of the EEG).

The subtraction spectrum EYES-OPEN-REST minus EYES-OPEN-MVT also exhibited a peak of alpha-band activity, with a topography maximal at central and bilateral parietal electrodes, compatible with activity originating from the hand representation within the left and right sensorimotor cortices (Figure 5B). As compared to the signals obtained using the ICA-based method, the contribution of 1/f noise was minimal in these subtracted spectra.

### Baseline peak alpha frequency (PAF) estimated using the CoG method

Using the CoG approach and a 9-11 Hz window applied to the VISUAL-ALPHA, SM-ALPHA and MIXED-ALPHA spectra obtained with the ICA-based method, the PAF of vision-related alpha-band activity estimated at occipital electrodes in the VISUAL-ALPHA signals during closure of the eyes was 10.0 ± 0.2 Hz (visual PAF). The PAF of sensorimotor-related alpha-band activity estimated at central electrodes in the eyes open condition at rest was 10.0 ± 0.1 Hz (sensorimotor PAF). The PAF estimated in the MIXED-ALPHA spectra was 10.0 ± 0.1 Hz at occipital electrodes during closure of the eyes, and 10.0 ± 0.1 Hz at central electrodes in the eyes open at rest condition. Using the same CoG approach applied to the spectra obtained with the subtraction-based approach, data from one participant was rejected for estimation of the visual PAF and data from three participants were rejected for estimation of the sensorimotor PAF as their spectra did not exhibit a peak within the frequency window. The visual PAF was 10.0 ± 0.3 Hz while the sensorimotor-related PAF was 9.9 ± 0.5 Hz. Average PAF estimates obtained using the larger 7-13 Hz frequency window are reported in Table 1.

**Table 1.**
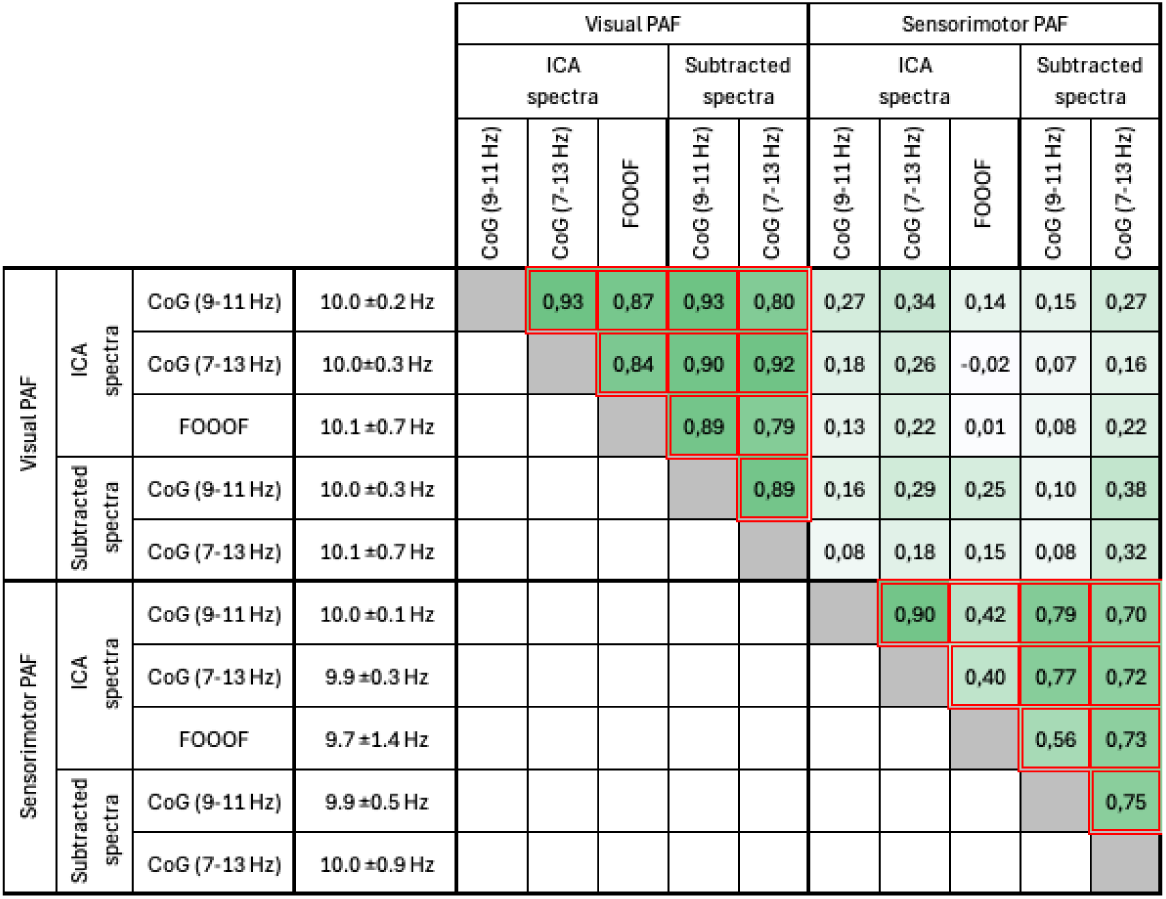
Spearman’s correlation coefficients between visual and sensorimotor PAF estimates obtained using the ICA-based or subtraction-based approaches to isolate visual- and sensorimotor-related alpha-band activity, and the CoG (frequency windows: 9-11 Hz and 7-13 Hz) or FOOOF methods to estimate peak frequency. Positive and negative correlations are coded in green and red, respectively. Significant correlations (p<0.05) are highlighted with red borders

### Estimation of Peak Alpha Frequency (PAF) using the FOOOF method

Using the FOOOF approach applied to the VISUAL-ALPHA, SM-ALPHA and MIXED-ALPHA spectra obtained with the ICA-based method, the PAF of vision-related alpha-band activity was 10.1 ± 0.7 Hz, while the PAF for sensorimotor-related alpha-band activity was 9.7 ± 1.4 Hz. The PAF estimated in the MIXED-ALPHA spectra was 10.3 ± 1.0 Hz at occipital electrodes during closure of the eyes, and 10.4 ± 1.2 Hz at central electrodes in the eyes open at rest condition. Average PAF estimates obtained using the larger 7-13 Hz frequency window are reported in Table 1.

### Primary analysis: correlation between baseline sensorimotor PAF and the area radius of HFS-induced secondary hyperalgesia

Using the data from the 24/32 participants in which the ICA-based approach succeeded in isolating sensorimotor-related alpha-band activity, no significant correlation between the sensorimotor PAF and the radius of the HFS-induced area of secondary hyperalgesia was found, neither when using the CoG estimates of PAF (CoG window 9-11 Hz: Spearman’s rho = 0.167, p = 0.434, Pearson’s r = 0.267, p = 0.207; CoG window 7 – 13 Hz: Spearman’s rho = 0.207, p = 0.330, Pearson’s r = 0.132, p = 0.539), nor when using the FOOOF estimates of PAF (Spearman’s rho = −0.203, p = 0.339, Pearson’s r = - 0.252, p = 0.235) (Table 2 and Figure 6).

**Figure 6.**
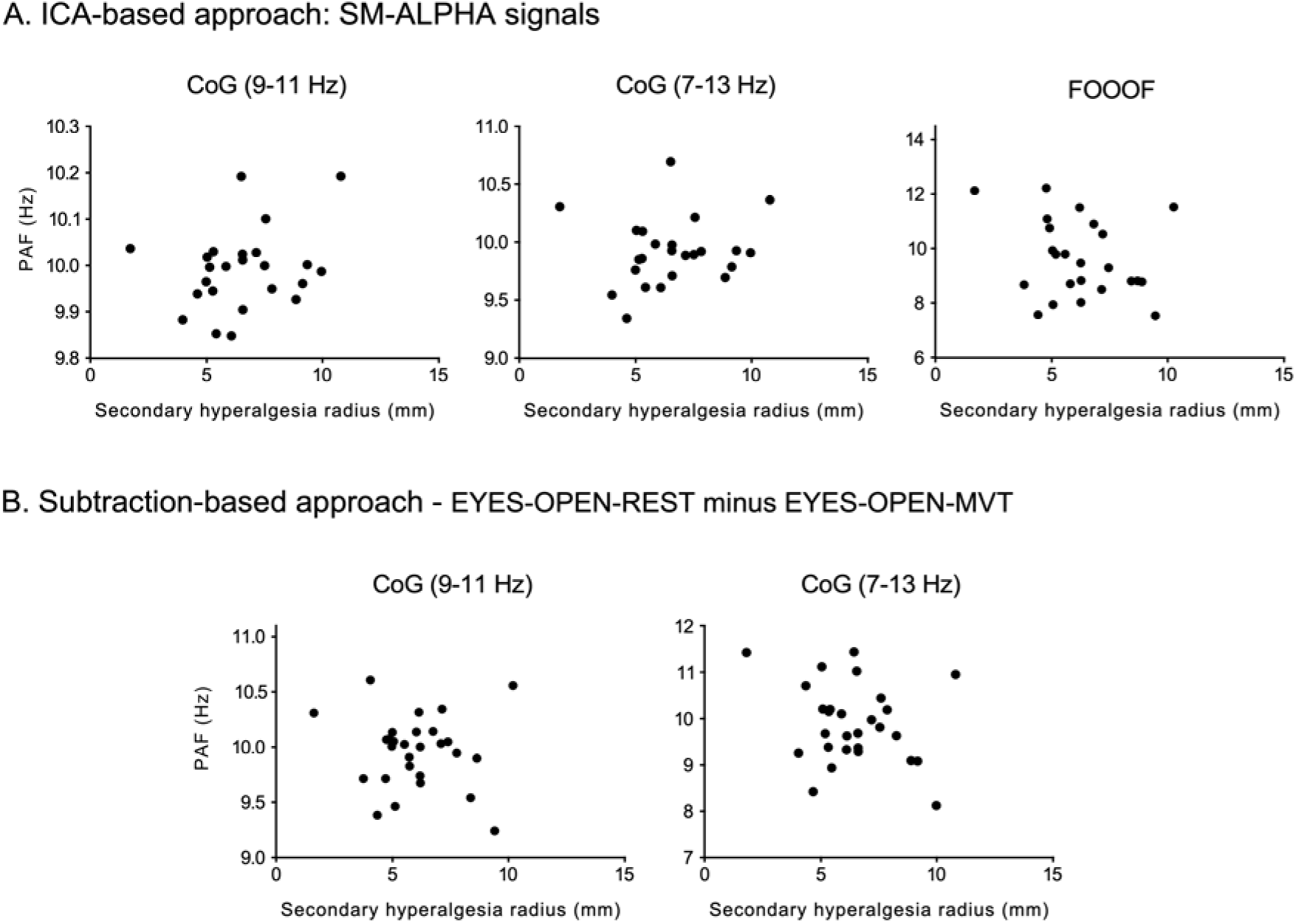
Scatter plots showing the absence of a significant Spearman’s correlation between the radius of the HFS-induced area of secondary hyperalgesia and the sensorimotor peak alpha frequency (PAF), regardless of the method used to isolate sensorimotor-related activity (A: ICA-based approach; B: subtraction-based approach) or the method used to estimate the PAF (center of gravity using a narrow 9-11 Hz or wide 7-13 Hz frequency window, FOOOF algorithm).

**Table 2.** Spearman’s correlation coefficients between visual and sensorimotor PAF estimates and the following measures: the area radius of HFS-induced secondary hyperalgesia area, the changes in pinprick sensitivity at the HFS-sensitized and non-sensitized forearms, the detection threshold to a single electrical pulse averaged across the two forearms, the baseline pinprick ratings averaged across the two forearms, and the pain ratings reported during HFS. Positive and negative correlations are coded in green and red, respectively. Significant correlations (p<0.05) are highlighted with red borders.

|  | Visual PAF |  |  |  |  | Sensorimotor PAF |  |  |  |  |
| --- | --- | --- | --- | --- | --- | --- | --- | --- | --- | --- |
|  | ICA spectra |  |  | Subtracted spectra |  | ICA spectra |  |  | Subtracted spectra |  |
|  | CoG (9-11 Hz) | CoG (7-13 Hz) | FOOOF | CoG (9-11 Hz) | CoG (7-13 Hz) | CoG (9-11 Hz) | CoG (7-13 Hz) | FOOOF | CoG (9-11 Hz) | CoG (7-13 Hz) |
| Secondary hyperalgesia radius | -0,23 | -0,33 | -0,23 | -0,12 | -0,28 | 0,17 | 0,21 | 0,20 | -0,07 | -0,10 |
| Change in pinprick sensitivity at HFS forearm ($\Delta\text{NRS}_{\text{HFS}}$) | -0,07 | -0,16 | 0,02 | -0,25 | -0,29 | -0,03 | -0,11 | -0,02 | 0,00 | -0,11 |
| Change in pinprick sensitivity at non-sensitized forearm ($\Delta\text{NRS}_{\text{NS}}$) | -0,49 | -0,42 | -0,44 | -0,48 | -0,36 | -0,17 | -0,25 | 0,00 | -0,01 | -0,08 |
| Change in pinprick sensitivity at HFS minus non-sensitized forearm ($\Delta\Delta\text{NRS}$) | 0,09 | -0,02 | 0,20 | -0,08 | -0,16 | 0,12 | 0,06 | -0,02 | 0,07 | -0,07 |
| Baseline pinprick ratings | 0,31 | 0,29 | 0,40 | 0,25 | 0,20 | 0,22 | 0,25 | -0,34 | -0,03 | 0,07 |
| HFS pain ratings | 0,20 | 0,09 | 0,31 | 0,15 | 0,01 | 0,36 | 0,39 | -0,03 | 0,15 | 0,17 |
| Electrical detection threshold | -0,29 | -0,11 | -0,17 | -0,12 | 0,03 | -0,13 | -0,12 | 0,05 | 0,11 | 0,05 |

Similarly, when using the PAF estimates obtained from the subtraction-based spectra, there was no significant correlation between the sensorimotor PAF and the radius of the HFS-induced area of secondary hyperalgesia (CoG window 9-11 Hz: Spearman’s rho=-0.068, p=0.729, Pearson’s r = −0.092, p = 0.641; CoG window 7-13 Hz: Spearman’s rho=-0.095, p=0.624, Pearson’s r = −0.107, p = 0.581) (Table 2 and Figure 6).

### Exploratory analyses

#### Correlation between baseline sensorimotor PAF and changes in pinprick sensitivity after HFS

Such as for the area of HFS-induced secondary hyperalgesia, there was no significant correlation between the sensorimotor PAF and (1) the magnitude of the HFS-induced increase in pinprick sensitivity at the sensitized forearm compared to the contralateral forearm (ΔΔNRS), (2) the change in pinprick sensitivity at the HFS-sensitized forearm (ΔNRS_HFS_) and (3) the change in pinprick sensitivity at the contralateral non-sensitized forearm (ΔNRS_NS_) (Table 2).

#### Correlation between baseline visual PAF and the area of HFS-induced secondary hyperalgesia

Such as for the sensorimotor PAF, there was no significant correlation between the visual PAF and the radius of the area of HFS-induced secondary hyperalgesia (Table 2).

#### Correlation between baseline visual PAF and the changes in pinprick sensitivity after HFS

There was also no significant correlation between the visual PAF and the increase in pinprick sensitivity at the sensitized forearm compared to the contralateral forearm (ΔΔNRS) or the change in pinprick sensitivity at the HFS-sensitized forearm (ΔNRS_HFS_). However, and regardless of the method used to estimate the visual PAF, there was a significant negative correlation between the visual PAF and the change in pinprick sensitivity at the non-sensitized forearm (ΔNRS_NS_): participants with a higher visual PAF at baseline exhibited a stronger perceptual habituation to pinprick stimulation at the non-sensitized forearm (Table 2 and Figure 7).

**Figure 7.**
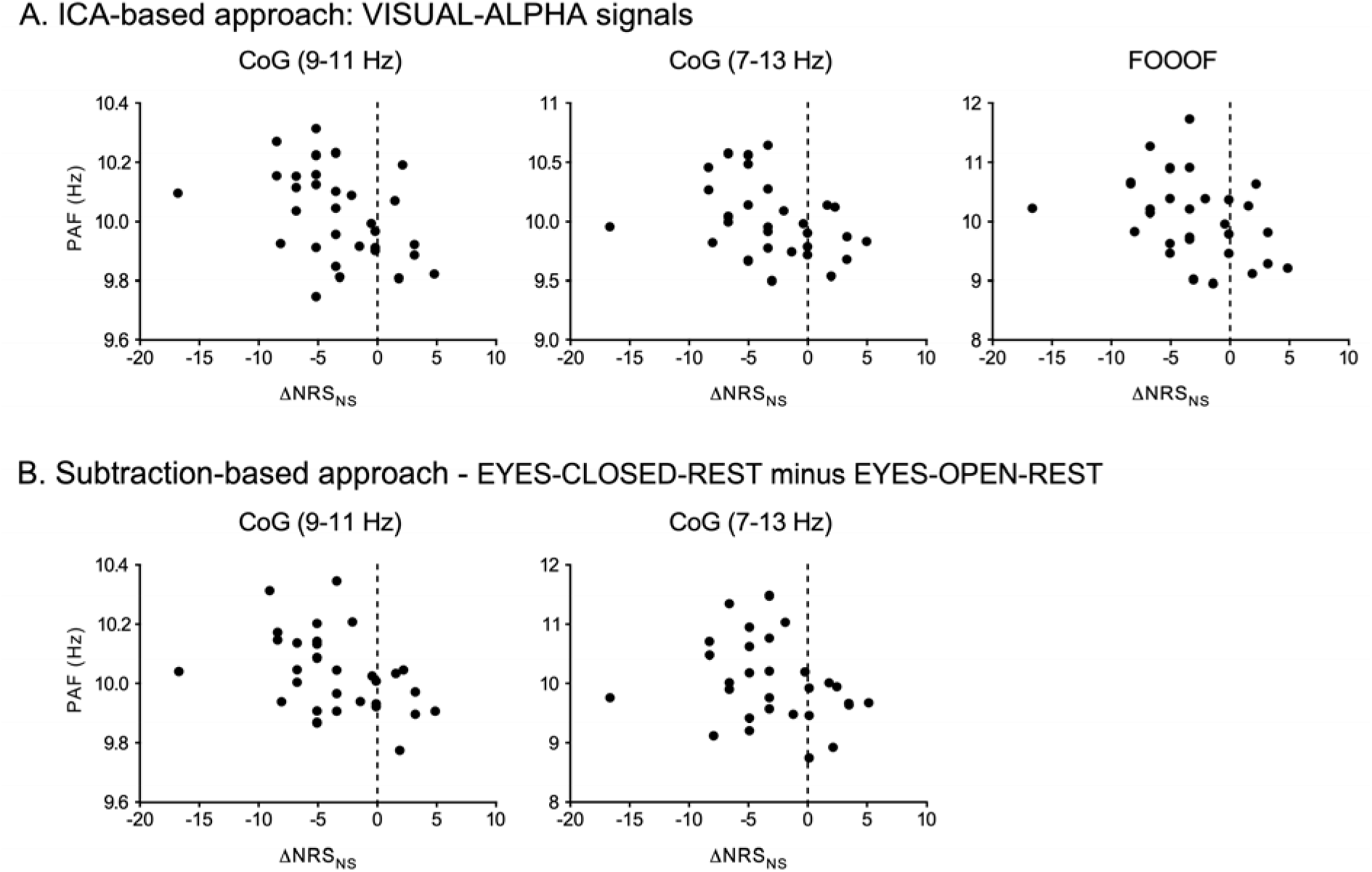
Scatter plots showing the Spearmans’s correlation between the post-HFS decrease in pinprick sensitivity at the non-sensitized forearm and the visual peak alpha frequency (PAF), present across methods used to isolate visual-related activity (A: ICA-based approach; B: subtraction-based approach) and methods to estimate the PAF (center of gravity using a narrow 9-11 Hz or wide 7-13 Hz frequency window, FOOOF algorithm).

#### Correlation between baseline PAF and baseline pain ratings

There was no significant correlation between either the visual PAF nor the sensorimotor PAF and (1) the detection threshold to single electrical pulses averaged across the two forearms, (2) the baseline pinprick ratings averaged across the two forearms, or (3) the intensity of the pain during HFS (Table 2).

#### Correlation between vision- and sensorimotor-related PAF

Both visual and sensorimotor PAF estimates derived using ICA-based and subtraction-based approaches to isolate visual- and sensorimotor-related alpha-band activity and computed using the CoG and FOOOF methods were highly correlated, indicating good consistency across the estimation techniques (Table 1). In contrast, there was no significant correlation between the visual PAF and the sensorimotor PAF, indicating that they likely reflect distinct oscillatory processes (Table 1). This highlights the importance of considering these rhythms as functionally separate when assessing their relevance to perception.

#### Mixed alpha-band activity

Using the ICA-based approach, independent components (ICs) capturing alpha-band activity that increased during closure of the eyes and decreased during finger movements were identified in 22/32 participants. The PAF estimates of these MIXED-ALPHA signals were strongly correlated with the visual PAF, but also showed significant correlation with the sensorimotor PAF using the largest window, indicating that these signals may reflect the fact ICA did not fully dissociate the two types of alpha-band activity (Table S1). There was no significant correlation between the PAF of MIXED-ALPHA signals and the secondary hyperalgesia radius, the changes in pinprick sensitivity after HFS, the electrical detection threshold and the HFS pain ratings. A weak correlation was reported for the baseline pinprick ratings and mixed PAF at occipital electrodes (Table S2).

## DISCUSSION

The results of the present study do not support our hypothesis that interindividual variations in peak-alpha frequency are associated with the susceptibility to develop HFS-induced secondary hyperalgesia. Indeed, in our sample of 32 healthy participants and using different approaches to isolate sensorimotor- and vision-related alpha-band activity and estimate peak frequency in the resting EEG, there was no significant correlation between sensorimotor or visual PAF estimates and neither the spatial extent of the HFS-induced secondary hyperalgesia area, nor the magnitude of the HFS-induced increase in pinprick sensitivity at the sensitized forearm.

### Functional dissociation between vision- and sensorimotor-related alpha-band activity

Previous reports of a relationship between PAF and sensitivity to pain have often interpreted the measured interindividual differences in PAF as reflecting differences in sensorimotor-related alpha-band activity. However, the topographical distribution of the alpha-band in resting EEG predominates over posterior areas, and its most prominent feature is its sensitivity to closure of the eyes. Hence, measures of PAF in resting EEG may be expected to mainly reflect vision-related alpha-band oscillations, even when the PAF is estimated at central electrodes overlying the sensorimotor cortex (17,19,55). For this reason, several authors have attempted to isolate sensorimotor-from vision-related alpha oscillations using an Independent Component Analysis and selecting components displaying a central topography as compared to an occipital topography (18,52,54,65).

In the present study, we propose two functional approaches to isolate vision- and sensorimotor-related alpha oscillations, based on the sensitivity of vision-related alpha oscillations to closure of the eyes and the sensitivity of sensorimotor-related alpha oscillations to movement execution rather than their topography. In the ICA-based approach, the functional properties of vision- and sensorimotor-related alpha-band activity allowed separating alpha-band activity into visual-related components that were sensitive to closure of the eyes and sensorimotor-related components that were sensitive to the performance of bilateral finger movements. In the subtraction-based approach, we subtracted the EEG spectra obtained during eyes open and eyes closed recordings at rest to isolate the alpha-band increase occurring during eye closure, while we subtracted the EEG spectra obtained during eyes open at rest and eyes open during bilateral finger movements to capture the change in alpha-band activity occurring during movement. Both the ICA-based approach and the subtraction-based approach showed that the alpha-band activity sensitive to closure of the eyes had a clear occipital topography compatible with activity originating from visual areas, whereas the alpha-band activity sensitive to moving of the fingers exhibited a bilateral central topography, compatible with activity originating from left and right sensorimotor areas.

It should be noted that in the ICA-based approach, visual-related alpha-band activity was isolated in 31/32 participants, whereas sensorimotor-related alpha-band activity was isolated in 24/32 participants. This may be related to the fact that the sensorimotor mu rhythm is not always observed (5).

### Visual and sensorimotor PAF estimates

Previous studies using the center-of-gravity (CoG) method to estimate PAF have shown that the choice of frequency window can markedly influence results (55,58,65). Some studies have employed relatively narrow windows (9-11 Hz) to minimize the influence of the aperiodic 1/f component of the EEG frequency spectrum (17–19,52), while other studies have used larger windows from 7 to 11 Hz (57), 7.4 to 12 Hz (56) or 7.5 to 13 Hz (58). In the present study, the CoG was computed using both narrow (9-11 Hz) and wide (7-13 Hz) windows to assess the robustness of the estimates. Additionally, CoG estimates computed on the subtracted spectra are expected to be minimally influenced by the 1/f aperiodic component, as this component should be largely cancelled by the subtraction procedure. Finally, we also used the FOOOF method to remove the aperiodic component and estimate PAF using a Gaussian fit rather than computing the center of gravity (62).

Both for the visual PAF and the sensorimotor PAF, there was a strong correlation between the different PAF estimates, indicating good consistency between estimation techniques.

Interestingly, and regardless of the method used to estimate PAF, the interindividual variations in visual PAF were not significantly correlated with the interindividual variations in sensorimotor PAF. This indicates that the two reflect at least partially distinct functional processes. Supporting this distinction, previous studies have shown that the mu-rhythm and alpha rhythm can differ in frequency and amplitude despite occupying the same frequency band (66). In light of these observations, future research should investigate whether the previously reported association between PAF and pain sensitivity (18,19,21,57,67) are predominantly driven by visual-versus sensorimotor-related alpha-band activity.

### Lack of association between PAF and the susceptibility to develop secondary hyperalgesia

Our study found no significant association between either visual or sensorimotor PAF and the susceptibility to develop HFS-induced secondary hyperalgesia. This suggests that the frequency characteristics of alpha-band activity do not play a modulatory role in the induction of central sensitization. Notably, recent studies examining whether cognitive factors that are known to modulate pain perception – such as attention and expectations – may also influence the susceptibility to develop experimentally-induced central sensitization have reported mostly negative results (68,69). Collectively, this suggests that the propensity to develop central sensitization may be minimally influenced by cognitive factors and the functional state of the brain.

Whether this absence of association between PAF and secondary hyperalgesia is specific for HFS-induced hyperalgesia in healthy volunteers, or whether it generalizes to other experimental methods to induce hyperalgesia, as well as to hyperalgesia in patients remains an open question requiring further studies.

Finally, it is important to acknowledge that the sample size in our study was based on prior work reporting relatively large correlation coefficients. As such, our study was underpowered to detect smaller effect sizes.

### Association between visual PAF and habituation at the non-sensitized forearm

An incidental finding of the present study was a negative association between visual PAF and perceptual habituation to pinprick stimulation. Specifically, participants with a higher visual PAF showed a more pronounced decrease in pinprick ratings at the non-sensitized forearm. This raises an intriguing possibility: could previous findings linking a lower PAF with greater pain sensitivity be partially explained by this relationship between PAF and habituation? If individuals with a lower PAF habituate less to repeated or sustained nociceptive stimuli as compared to individuals with a higher PAF, they would also be expected to report higher average pain ratings when exposed to such stimuli. Such an interpretation would also be compatible with the fact that the studies having shown a relationship between PAF and sensitivity to pain have relied on methods inducing sustained pain such as capsaicin-heat pain, NGF injection, and tonic heat pain (18–20,67), while studies employing more phasic pain stimuli have reported no or only marginally-significant associations between PAF and pain ratings (70).

## CONCLUSION

In this study conducted in healthy volunteers, interindividual variations in visual or sensorimotor PAF were not significantly associated with differences in the susceptibility to develop HFS-induced secondary hyperalgesia. This suggests that PAF does not modulate the propensity to develop central sensitization. However, exploratory analyses revealed that individuals with a higher visual PAF tended to exhibit stronger perceptual habituation to pinprick stimulation. Further studies are needed to confirm this incidental observation, and examine whether it might contribute to the previously reported association between PAF and pain sensitivity.

## DATA AVAILABILITY

Source data are available at https://doi.org/10.6084/m9.figshare.29145725

## SUPPLEMENTAL MATERIAL

Supplemental Fig. S1 & Table S1-S2 DOI: https://doi.org/10.6084/m9.figshare.29145695

## CURRENT ADDRESS

Same as affiliations

## DISCLAIMERS

The statements and opinions presented here reflect the author’s view and neither IMI nor the European Union, EFPIA, or any Associated Partners are responsible for any use that may be made of the information contained therein.

## AUTHORS CONTRIBUTIONS

Louisien Lebrun^1^*, Gloria Ricci^2^, Arthur Courtin^1,3^, Emanuel N. van den Broeke^1,4^, Cédric Lenoir^1^, André Mouraux^1^

Louisien Lebrun: conceived and designed research, performed experiments, analyzed data, interpreted results of experiment, prepared figures, drafted manuscript, approved final version of manuscript

Gloria Ricci: conceived and designed research, analyzed data, edited and revised the manuscript, approved final version of manuscript

Arthur Courtin: conceived and designed research, edited and revised the manuscript, approved final version of manuscript

Emanuel N. van den Broeke: conceived and designed research, edited and revised the manuscript, approved final version of manuscript

Cédric Lenoir: conceived and designed research, edited and revised the manuscript, approved final version of manuscript

André Mouraux: conceived and designed research, analyzed data, interpreted results of experiment, prepared figures, edited and revised the manuscript, approved final version of manuscript

