## Supplemental Fig. S1 & Table S1-S2 for "Interindividual variations in peak alpha frequency do not predict the magnitude or extent of secondary hyperalgesia induced by high-frequency stimulation"

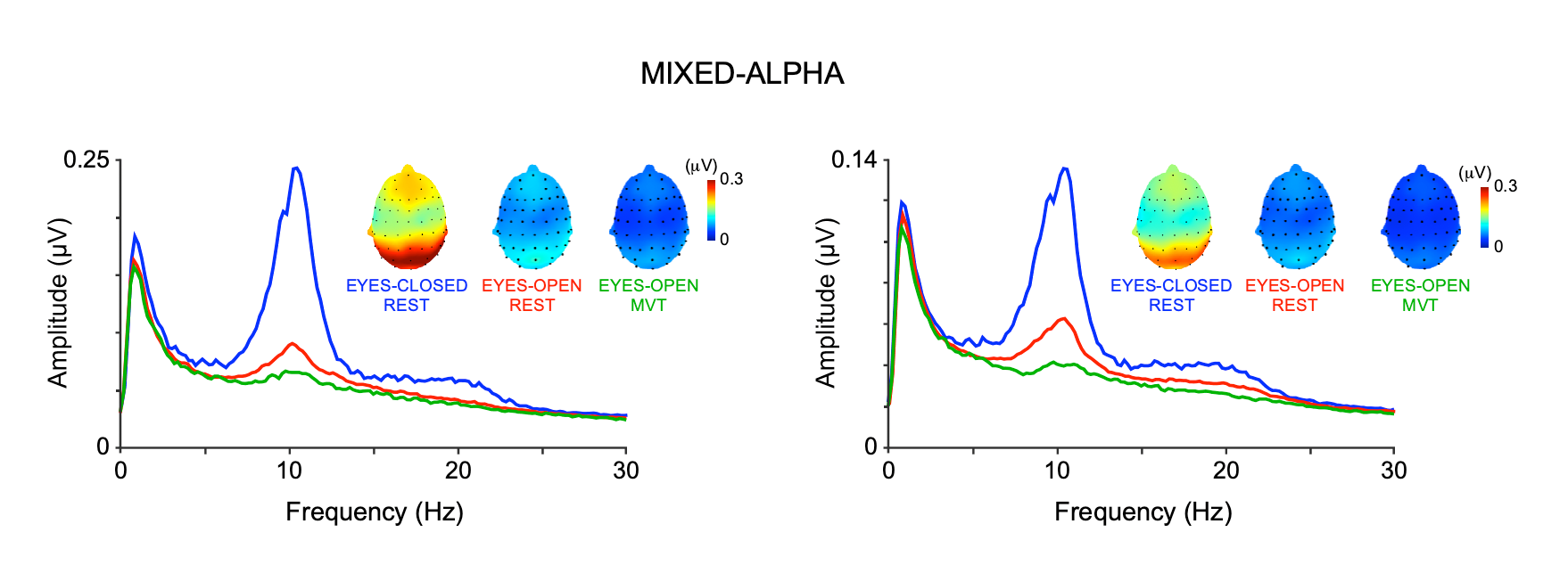

**Figure S1**. Group-level average frequency spectra of the ICA-separated signals capturing alpha-band activity that both increased during closure of the eyes, and decreased during movement of the fingers (MIXED-ALPHA signals) at occipital (left) and central (right) electrodes. Note that the alpha-band activity was maximal over occipital regions, compatible with activity originating predominantly from visual areas, both when computing the alpha-peak topography at occipital electrodes and at central electrodes.

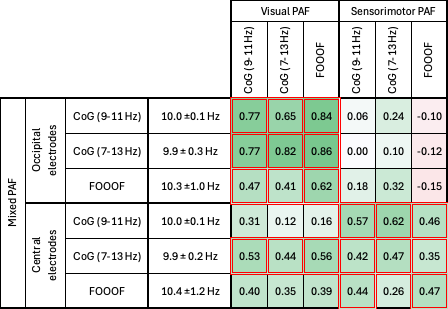

**Table S1**. Spearman’s correlation coefficients between the PAF of mixed alpha-band activity (activity that was sensitive to both closure of the eyes and bilateral finger movements, PAF assessed at occipital and central electrodes) and the visual and sensorimotor PAF, using the estimates obtained using the ICA-based or subtraction-based approaches to isolate visual- and sensorimotor-related alpha-band activity, and the CoG (frequency windows: 9-11 Hz and 7-13 Hz) or FOOOF methods to estimate peak frequency. Positive and negative correlations are coded in green and red, respectively. Significant correlations (p<.05) are highlighted with red borders

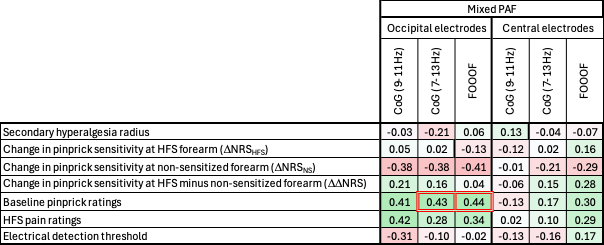

**Table S2**. Spearman’s correlation coefficients between the PAF of mixed alpha-band activity (activity that was sensitive to both closure of the eyes and bilateral finger movements, PAF assessed at occipital and central electrodes) and the following measures: the area radius of HFS-induced secondary hyperalgesia area, the changes in pinprick sensitivity at the HFS-sensitized and non-sensitized forearms, the detection threshold to a single electrical pulse averaged across the two forearms, the baseline pinprick ratings averaged across the two forearms, and the pain ratings reported during HFS. Positive and negative correlations are coded in green and red, respectively. Significant correlations (p<.05) are highlighted with red borders.
